# IFI207, a young and fast-evolving protein, controls retroviral replication via the STING pathway

**DOI:** 10.1101/2024.04.30.591891

**Authors:** Eileen A. Moran, Karen Salas-Briceno, Wenming Zhao, Takuji Enya, Alexya N. Aguilera, Ivan Acosta, Francis Alonzo, Dara Kiani, Judith Behnsen, Catalina Alvarez, Thomas M. Keane, David J. Adams, Jingtao Lilue, Susan R. Ross

## Abstract

Mammalian AIM-2-like receptor (ALR) proteins bind nucleic acids and initiate production of type I interferons or inflammasome assembly, thereby contributing to host innate immunity. In mice, the *Alr* locus is highly polymorphic at the sequence and copy number level and we show here, is one of the most dynamic regions of the genome. One rapidly evolving gene within this region, *Ifi207*, was introduced to the *Mus* genome by gene conversion or an unequal recombination event a few million years ago. *Ifi207* has a large, distinctive repeat region that differs in sequence and length among *Mus* species and even closely related inbred *Mus musculus* strains. We show that IFI207 controls MLV infection *in vivo* and that it plays a role in the STING-mediated response to cGAMP, dsDNA, DMXXA and MLV. IFI207 binds to STING and inclusion of its repeat region appears to stabilize STING protein. The *Alr* locus and *Ifi207* provide a clear example of the evolutionary innovation of gene function, possibly as a result of host-pathogen co-evolution.

**IMPORTANCE:** The Red Queen hypothesis predicts that the arms race between pathogens and the host may accelerate evolution of both sides, and therefore cause higher diversity in virulence factors and immune-related proteins, respectively (1). The *Alr* gene family in mice has undergone rapid evolution in the last few million years and includes the creation of two novel members, *MndaL* and *Ifi207*. *Ifi207* in particular became highly divergent, with significant genetic changes between highly related inbred mice. IFI207 protein acts in the STING pathway and contributes to anti-retroviral resistance via a novel mechanism. The data show that under the pressure of host-pathogen coevolution in a dynamic locus, gene conversion and recombination between gene family members creates new genes with novel and essential functions that play diverse roles in biological processes.

## Introduction

Innate immune responses are critical to control viral, bacterial, and fungal infections. The mammalian innate immune system consists of numerous pattern recognition receptors (PRRs) that recognize intracellular nucleic acids and other components of pathogens, termed pathogen-associated molecular patterns (PAMPs) (2). PRR activation initiates intricate signaling cascades which culminate in the production of type I interferons (IFNs), proinflammatory cytokines, and inflammatory cell death. These responses also ensure that the adaptive immune system is activated and subsequently generates a specific response to eliminate pathogens. Multiple PRRs likely evolved so that the innate immune system can discriminate between the many different pathogens and PAMPs it encounters (3–6).

Many anti-pathogen factors show high levels of genetic diversity between species with regard to both sequence and copy number, including type I interferons, apolipoprotein B mRNA editing enzyme catalytic polypeptide 3 (APOBEC3) and Tripartite Motif Containing 5 (TRIM5) (7–9). Another example of this is the AIM2-like receptor (ALR) family; the number of *ALR* genes varies from 4 in humans to 13-16 in mice, all encoded at a single locus (10–13). Although it is not known what selective pressures caused the diversification of this locus, several ALRs have been studied for their role in innate immunity. Human interferon-gamma inducible (IFI) 16 and mouse IFI204 are required for type I IFN and cytokine responses to herpes simplex virus-1 (HSV-1) and vaccinia virus (VACV) DNA (14). Several ALRs also affect transcription; IFI16 and IFI204 function in the nucleus to regulate HIV and HSV-1 transcription, while IFI205 regulates the inflammatory response by controlling apoptosis-associated speck-like molecule containing CARD domain (ASC) transcription (15–17) As cytoplasmic sensors, IFI204 is required for type I IFN production during *M. bovis* and *F. novicida* infection, and IFI203 and IFI205 play a role in the IFN response to murine leukemia virus (MLV) infection and endogenous retroelement DNA detection, respectively (12, 18–20). These ALRs, as well as the innate immune sensors like cyclic GMP-AMP (cGAMP) synthase (cGAS) and DEAD-Box Helicase 41 (DDX41), mediate production of IFNs and cytokines in a stimulator of interferon genes (STING)-dependent manner (4, 12, 17–23). Activated STING dimerizes and interacts with TANK binding kinase 1 (TBK1), thereby stimulating the dimerization and nuclear translocation of the transcription factors IFN regulatory factor 3 (IRF3) and NF-κB, leading to the production of type I IFNs and cytokines (24–27).

Here we show that the *Alr* locus in *Mus* species has undergone rapid evolution for at least the past few million years, and that one gene in the locus, *Ifi207* (*PyhinA*), is highly divergent even among closely related inbred mouse strains. We found that IFI207 is unique among the ALRs due to a large repeat region separating the two domains common to all ALRs, the N-terminal pyrin domain (PYD) that facilitates homotypic and heterotypic interactions with PYD-containing and other proteins, and the C-terminal hematopoietic expression, interferon-inducible nature, and nuclear localization (HIN) 200 domain that binds DNA (5, 10, 28). We found that the number of repeats is variable among inbred wild and laboratory strains of mice, ranging in number from 11 to 25. Due to the evidence of pathogen-driven positive selection on the *Alr* locus and the evolutionary diversity of *Ifi207*, we examined the role of IFI207 in innate immune responses. We demonstrate that IFI207 binds to STING protein and stabilizes it via its repeat region. This binding enhances the STING response and contributes to the control of MLV infection *in vivo*.

## RESULTS

### *Ifi207* is a young gene exclusively found in the *Mus* genus

We previously reported that the *Alr* locus is highly variable, with different numbers and complements of *Alr* genes mapping to the locus in three inbred mouse strains, C57BL/6, DBA/2J and 129P2/OlaHsd (12). The locus in the GRCm39 reference genome (strain C57BL/6J) encodes 14 genes (Fig. 1A), whilst strain 129P2/OlaHsd encodes 16, and DBA/2J, 17 genes (12). Many members of the *Alr* family are similar to each other, both in the coding and non-coding regions, and are surrounded by transposon elements. To further study the genome structure of *Alr* locus in house mice, we accessed the long read (whole genome PacBio Sequel / Hifi + Chicago HiC and BioNano scaffolding) *de novo* assemblies of 13 inbred house mouse strains, *Mus Spretus* and *Mus Caroli* from the Mouse Genomes Project; the genome assemblies will be described in a separate manuscript. As shown in Fig. 1A and 1B, based on gene structure and phylogeny, the *Alr* gene family consists of 9 variant members in house mice and other *Mus* species, and the number of genes in the locus ranges from 10 (PWK/PhJ) to 17 (DBA/2J, AKR/J, CBA/J) (Fig. 1A). In inbred mouse strains, there is one copy of *Aim2* at the 5’ end of the locus followed by 2-3 copies of *Ifi206* homologues (*Ifi206*, *Ifi213* and *Ifi208* in the GRCm39 reference), 1-2 copies of *Ifi214*-like genes (*Ifi214* and *Ifi209*), one copy of *Ifi207*, 1-3 copies of *Ifi202*, 1-4 *Ifi203* homologues/alleles, 1-2 *Mnda* members (*Ifi205* or *Mnda*), one copy of *Ifi204* and possibly one copy of *MndaL*. *Ifi208*, *Ifi207*, *Ifi204* and one copy of *Ifi203* are encoded in a stable island in the middle of *Alr* locus, whilst other regions are highly dynamic in the evolutionary history of the house mouse.

**FIG 1.**
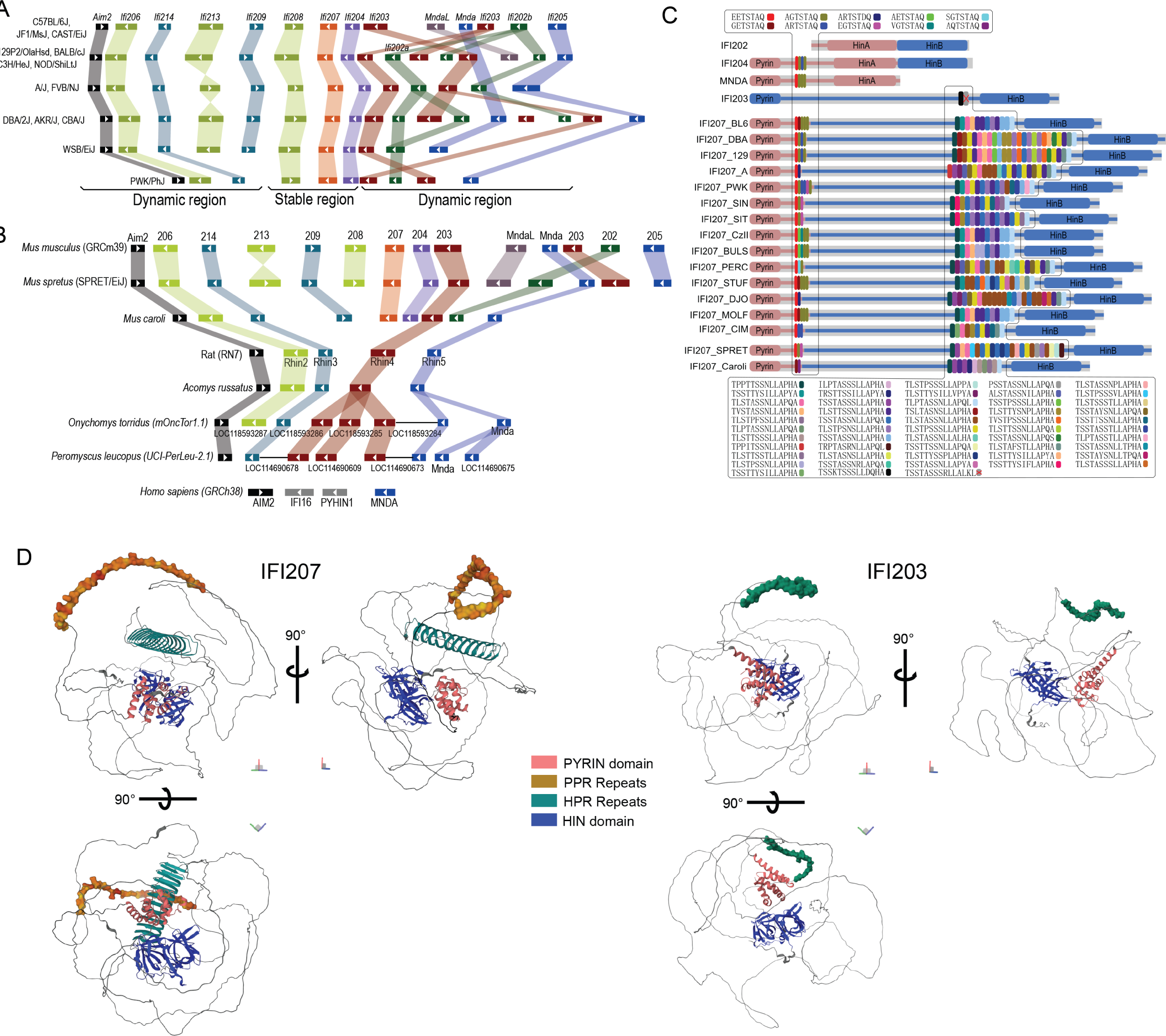
*Ifi207* structure. A) Genome structure of the *Alr* locus in inbred mouse strains. Based on gene structure and phylogeny, the 9 family members with variant copy numbers are indicated by the color blocks. *Ifi208*, *Ifi207*, *Ifi204* and one copy of *Ifi203* are encoded in a stable contig, and other regions in the *Alr* locus showed significant numbers of inversions, recombinations and duplications. B) Genome structure of the *Alr* locus in other rodent species and humans. *Ifi207*, *MndaL* and *Ifi202* are only found in the *Mus* genus. Some of the genes in *Onychomys* and *Peromyscus* were annotated tandemly, probably caused by low annotation quality. C) IFI207 protein structure in different *Mus* genus members. Shown for comparison is the structure of IFI202, IFI204, MNDA and IFI203 (see Tables S2 and S3 for sequences). Similar PYRIN domains are indicated by color, as are the HIN-A and HIN-B domains. The long linker sequence between these domains in IFI203 and IFI207 is indicated by a blue bar. D) Comparison of C57BL/6 IFI207 (E9Q3L4) and IFI203 (E9QAN9) predicted protein structures. Structures were downloaded from AlphaFold (https://alphafold.ebi.ac.uk/). Domains are colored as follows: PYD domain (pink), PYD-proximal repeat (PYD_PR_) (brown), HIN-proximal repeat (HIN_PR_) (teal), HIN-B (blue).

In addition, we downloaded the whole genome assembly from similar long read assemblies carried out with other rodent species. When the *Mus* locus is compared with other rodent species, only orthologues of *Aim2*, *Ifi206*, *Ifi214*, *Mnda* and *Ifi203* can clearly be found in the latter (Fig. 1B). All rat assemblies available on NCBI (*<u>Rattus norvegicus</u>*, strain BN/NhsdMcwi, SHRSP/BbbUtx, WKY/Bbb, SHR/Utx, SS/JrNhsd and GK/Ox; *Rattus rattus* CSIRO assembly) contain exactly 5 *Alr* members. The Golden spiny mouse (genus *Acomys,* subfamily *Deomyinae*), deer mice (genus *Peromyscus*) and grasshopper mice (*Onychomys*; both from family *Cricetidae*) have the same genome structure as rats, except with variable copy numbers of *Ifi203* and *Mnda* paralogues. No *Ifi207*, *MndaL*, *Ifi204* and *Ifi202* orthologues can be found in these species (Fig. 1B). Furthermore, we compared the *de novo* assembly of 29 species in the *glires* clade (rodents and rabbits) and investigated the whole genome Illumina reads of key species in the *Murinae* subfamily. Homologues of *Ifi207* can only be found in the *Mus* genus and arguably in the species *Apodemus sylvaticus* (Figs. S1 and S2 and Table S1; see Discussion). Indeed, the genome structure around *Ifi207* (including introns) is probably derived from the 3’ end of *Ifi203* and the 5′ end of *Mnda*, from a recent recombination event in the ancestor of the *Mus* genus. Additionally, the 5’ end of *Ifi203* and the 3’ end from *Mnda* possibly fused into *MndaL* in the same event (Fig. S3). In comparison to rodent species, the human genome encodes *AIM2* and *MNDA*, and two other genes with ambiguous relationship to rodent *Alrs*, namely *IFI16* and *PYHIN1* (Fig. 1B) (11).

### *Ifi207* is characterized by two highly polymorphic repeat regions

Most ALRs contain an N-terminal PYD domain and one or two HIN domains, termed HIN-A and HIN-B, depending on their sequence (10, 11). Comparison of the amino acid sequences of IFI204, MNDA and IFI207 demonstrated high homology in the PYRIN domain (Fig. 1C). Moreover, IFI203 and IFI207 share a long linker sequence with 72% identity between the PYRIN and HIN-B domains (Fig. 1C; Tables S2 and S3).

From the *de novo* assembly of the *Alr* locus in mice, we found two repeat regions in several genes which are highly polymorphic between mouse strains. At around residue 120 (PYD-proximal repeat) of IFI204, IFI205, MNDA and IFI207 there are 2-6 units of the sequence XXTSTA/DQ. All the repeats are encoded in the same exon, precisely 7 residues as a unit, and the first two residues are highly divergent (A, E, G, R, V, S or Q; Fig. 1C). This repeat was also found in the 2nd exon of *Mnda* in other rodent species, but not in humans (Fig. S1, Tables S2 and S3).

The other repeat unit is unique to IFI207 and is located close to the HIN-B domain, with 11 - 25 units of the sequence TXSTXSSXLLAPXA (S/T-rich repeat). In 49 unique repeat sequences extracted from 16 distinct alleles (extracted from 15 Mouse Genome Project (MGP) sequences plus 9 wild-derived mouse strains), there was up to 36% diversity (average value 15%) between their nucleotide sequence, and up to 65% difference at the amino acid residue level (29% average); this was confirmed with multiple sequencing methods (Fig. 1C and Tables S2 and S3; see Materials and Methods). Like the PYD-side repeat, all the S/T-rich HIN-side repeats are encoded in a single large exon, with precisely 14 residues (42 base pairs) in a unit. Interestingly, in IFI203, the possible origin and closest paralogue of IFI207, the sequence surrounding the repeat region is conserved between IFI203 and IFI207; however, in IFI203 only one or two copies of the S/T-rich HIN-side repeats are found, and they are highly conserved between mouse strains (Fig. S2). Moreover, a single copy of the same S/T-rich sequence can also be found close to the HIN-B domain of human IFI16 (Fig. S2, Table S2). These results indicate the IFI207 long S/T-rich repeats were recently introduced and are evolving in a very short time in mouse evolution, approximately 6.5 million years ago in an ancestor of the *Mus* genus (29) (see Table S3 for the alignment of Ifi203 and Ifi207).

AlphaFold analysis predicts that the S/T-rich repeat region forms a coiled structure not found in the other proteins encoded in the *ALR* locus; shown for comparison is the predicted structure of IFI203, the most closely related ALR (Fig. 1D). IFI207 from *Mus caroli* is also predicted to have a similar coiled structure (Fig. S4). The extremely fast evolution of the S/T-rich repeats in *Ifi207* make it one of the most dynamic regions in the mouse genome, and indeed, in mammalian species in general.

### Knockout of *Ifi202b*, *Ifi204* or *Ifi207* in mice has a minimal effect on expression of other genes in the locus

Mouse *Alr* genes, including both coding and intervening sequences, are highly homologous, making it difficult to precisely delete single genes in the locus (12). However, we were able to use CRISPR/Cas9 to delete exon 3 of the *Ifi207* gene in C57BL/6N mice (Fig. S5A). Additionally, mice with floxed alleles of *Ifi202b* and *Ifi204* on a C57BL/6N background were available through the KOMP repository. *Ifi202b* encodes a protein that lacks a PYD and has two HIN domains, A and B, while *Ifi204* encodes a protein with PYD, HIN-A and HIN-B domains (Fig. 1C). The replacement allele in IFI202 KO mice deleted exon 3, and Cre-mediated deletion resulted in IFI204 KO mice lacking exons 3 and 4 (Fig. S5A; see Methods). The knockout alleles for both *Ifi204* and *Ifi207* generate truncated proteins predicted to contain only the PYD, while the *Ifi202* knockout allele generates a protein predicted to contain a portion of the HIN-A domain and a full HIN-B domain (Fig. S5B). All three KO strains had normal viability and produced offspring.

We examined expression of the KO alleles in bone marrow-derived macrophages (BMDMs) and dendritic cells (BMDCs) from wild type (WT; BL/6) and knockout mice using reverse transcription quantitative PCR (RT-qPCR). *Ifi202* RNA was barely detectable in both cell types from WT mice, as we previously reported (12); this was also the case in the KO mice (Fig. S6A). *Ifi204* and *Ifi207* RNAs were not detected in the respective KO strains (Fig. S6A).

To determine if *Ifi202*, *Ifi204* or *Ifi207* knockout affected expression of the other genes in the locus, we examined RNA levels in BMDMs and BMDCs from WT, IFI202, IFI204 and IFI207 KO mice. BMDMs from STING^mut^ mice were also included in these analyses. These mice are resistant to stimulation with DNA and other PAMPs as they encode a mutant STING that is unable to signal (30). STING RNA levels were equivalent in WT, IFI207 KO and STING^mut^ BMDMs (Fig. S6B). Because it has previously been shown that the genes in the *Alr* locus are type 1 IFN-inducible, we also tested whether deletion of *Ifi202*, *Ifi204* or *Ifi207* affected induction of other *Alrs* by IFNβ (11, 31, 32). Neither basal nor IFNβ-induced levels of expression of the different genes were altered in either BMDMs or BMDCs by knockout of any of the ALRs or the STING mutation, apart from *Mnda*; *Mnda* RNA levels were consistently reduced in BMDMs and BMDCs in IFI204 KO mice in the absence and presence of IFNβ (Fig. S6A and 6B). The significance of this observation is unknown. Nonetheless, since loss of *Ifi207* did not affect the expression of the other *Alrs*, phenotypes observed in the KO mouse are likely specific to this gene.

### IFI207 is an antiviral restriction factor in mice but does not affect the cytokine response to bacteria

We showed previously that multiple activators of the STING pathway, including cGAS and DDX41, were needed to control *in vivo* MLV infection (18, 21, 33). To determine if IFI207 played a role in controlling virus infection, we infected newborn IFI207 and IFI204 KO mice, as well as mice heterozygous for the *Ifi207* KO allele with MLV. STING^mut^ mice, which we showed previously are highly susceptible to MLV infection, were infected for comparison (18, 21). Splenic viral titers were measured at 16 dpi. MLV titers were significantly higher in both IFI207 KO and heterozygous mice compared to WT mice (Fig. 2A). Similar results were seen in mice infected mice with a mutant virus, MLV^gGag^ which has an unstable capsid; overall titers were lower than in wild type MLV-infected mice, which we showed previously was due to the increased susceptibility of MLV^gGag^ to nucleic acid sensors and APOBEC3-mediated restriction (33). STING^mut^ mice were more highly infected with both wild type MLV and MLV^gGag^ compared to IFI207 KOs, like our previous studies with knockout of cGAS and DDX41, upstream effectors of STING (18, 21). IFI207 heterozygous mice, which express half as much *Ifi207* RNA compared to WT mice, also showed higher levels of infection than WT mice, suggesting haploinsufficiency (Fig. 2A). Finally, although it has been suggested that IFI204 is an anti-retroviral restriction factor, IFI204 KO mice had titers equivalent to WT mice (17). These data demonstrate that IFI207 is one of several factors that control MLV infection *in vivo*.

**FIG 2.**
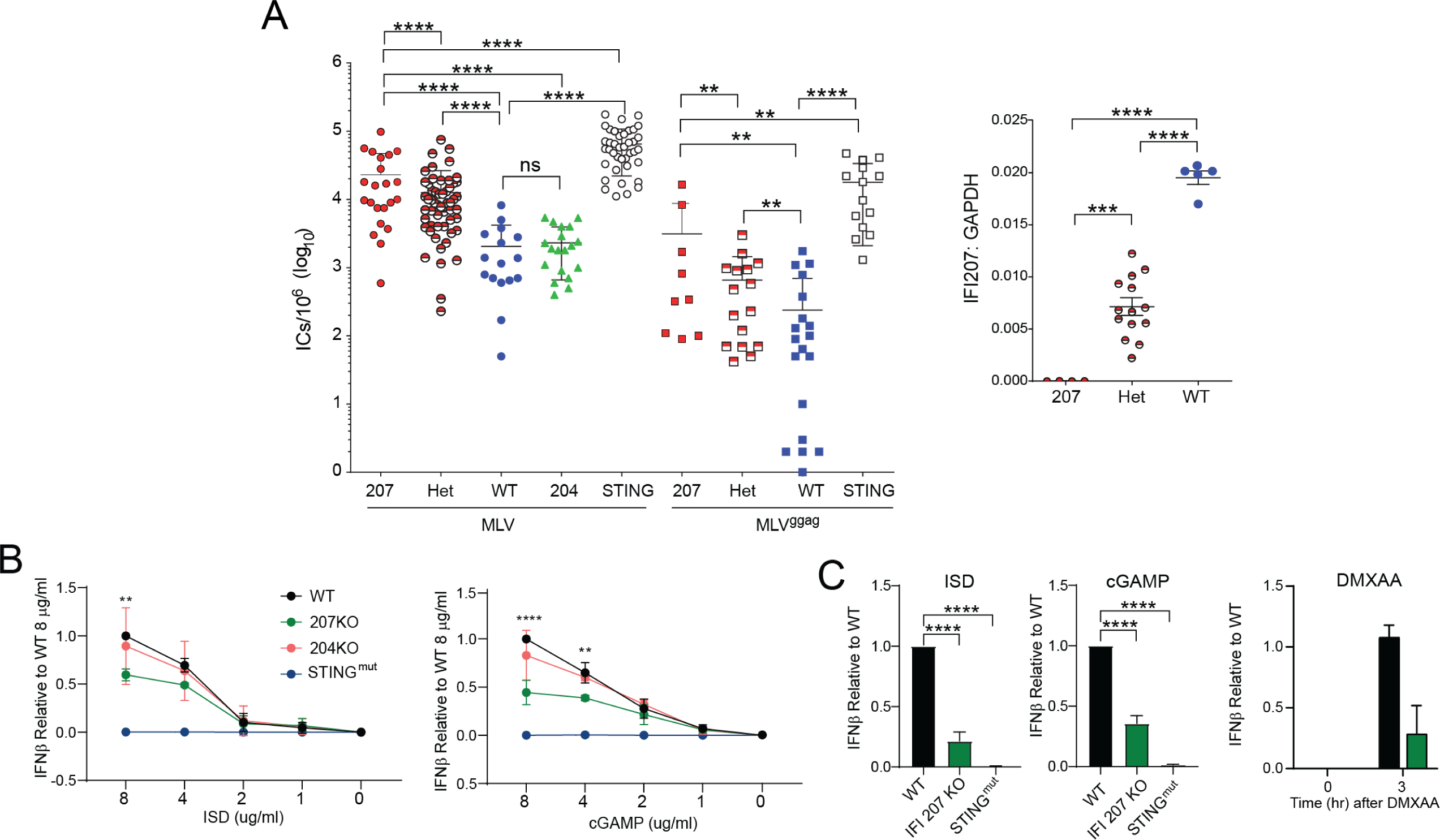
IFI207 KO mice are more susceptible to MLV infection. (A) Pups of the indicated genotype were infected with 2000 pfu of MLV (MLV or MLV^gGag^). At 16 dpi, splenic viral titers were determined on NIH3T3 cells by infectious center (IC) assays. Significance was determined by two-tail Mann-Whitney test (**, P<0.003; ****, P<0.0001). Right panel: RT-qPCR analysis to verify *Ifi207* RNA levels in mice. Symbols represent individual mice. Significance was determined by one-way ANOVA (***, P<0.002; ****, P<0.0001). (B) BMDMs from the indicated mice were transfected with the indicated concentrations of ISD or cGAMP for 3 hr. *Ifnβ* RNA levels were quantified by RT-qPCR. Shown are the averages of 3 independent experiments. (C) Primary fibroblasts from IFI207 KO and STING^mut^ mice were transfected with 8μg/ml of ISD or cGAMP or 100μg/ml DMXAA and at 3 hr post-transfection *Ifnβ* RNA levels were measured. Shown is the average of 3 experiments (ISD and cGAMP) or 2 experiments (DMXAA). Significance was determined by two-way ANOVA (****, P<0.0001)

A recent publication suggested that *Ifi207*-depleted cells had a diminished TNFα and IFNβ response to bacterial infection because of the loss of RNAPolII at the promoters of the genes (34). We also tested the response of BMDMs from IFI207 KOs to *S. aureus* and *K. pneumoniae*. *Tnf* and *Ifnβ1* RNA levels were similar in WT, IFI204 and IFI207 KO BMDMs in response to *S. aureus*. *Ifnβ1* but not *Tnf* expression was reduced in STING^mut^ cells, as previously reported (Fig. S7A) (35). In contrast, there was no significant difference in the *Tnf* or *Ifnβ1* response to *K. pneumoniae* in any of the mutant cells compared to WT (Fig. S7B).

### IFI207 plays a role in STING pathway signaling

Type I interferons generated by activation of the STING pathway are needed to control *in vivo* infection by MLV (18, 21). Several innate immune pathways can be activated to induce the production of interferon, including the TOLL-like receptor (TLR), RIG-I/MDA5 and STING pathways. To determine which pathway might be affected by loss of *Ifi207*, we used immortalized NR9456 macrophages in which *Ifi207* levels were reduced by siRNAs, as well as primary BMDMs from WT, and IFI207 and TLR4 KO mice. In both NR9456 cells and BMDMs, *Ifi207* depletion did not affect LPS induction of *Tnfα* or *Ifnβ1* RNA (Figs. S8A and 8B). In contrast, induction of both cytokine RNAs was ablated in BMDMs isolated from TLR4 knockout mice (Fig. S8B). As expected, since *Ifi207* is itself type I IFN-inducible, LPS treatment increased *Ifi207* levels in the immortalized macrophages, although to a lesser extent in the siRNA-depleted cells (Fig. S8A). LPS-treated WT but not IFI207 KO BMDMs had high levels of *Ifi207* RNA (Fig. S6B). Similar results were seen *in vivo* when LPS was subcutaneously injected; WT and IFI207 but not TLR4 KO mice had increased *Tnf* RNA in the draining lymph nodes (Fig. S8C).

We then examined whether IFI207 altered STING-mediated type I IFN induction. BMDMs from WT, IFI207 KO, IFI204 KO and STING^mut^ mice were transfected with increasing amounts of interferon stimulatory DNA (ISD) or cGAMP, the activating ligand of STING. Three hours (hr) after transfection, *Ifnβ1* RNA levels were measured. The IFN response to ISD and cGAMP was significantly reduced in IFI207 KO BMDMs, while the response by WT and IFI204 KO cells was the same; STING^mut^ cells had no response (Fig. 2B). Similar results were obtained when fibroblasts from WT, IFI207 KO and STING^mut^ were transfected with ISD or cGAMP (Fig. 2C). The IFN response to the STING agonist 5,6-Dimethylxanthenone-4-acetic acid (DMXAA) was also decreased in IFI207 KO fibroblasts (Fig. 2C). Thus, the presence of IFI207 in multiple cell types enhanced the STING response.

### STING protein levels and signaling are decreased in the absence of IFI207

IFI207 could be acting as an upstream sensor to activate STING, or it could be directly interacting with it to enhance responses. First, we determined if reducing endogenous IFI207 levels altered basal levels of STING. Transfection of IFI207 siRNAs into in NIH3T3 cells reduced basal STING levels compared to control siRNA-transfected or untransfected cells (Fig. 3A). Similar results were seen in IFI207 KO macrophages (Figs. 3B and 3D), fibroblasts (Fig. 3C) and DCs (Fig. S9A) (untreated lanes). Although IFI207 KO cells had lower levels of STING protein, they had similar levels of STING RNA to WT or STING^mut^ cells, demonstrating that IFI207 was not altering STING transcription (Fig. S5B).

**FIG 3.**
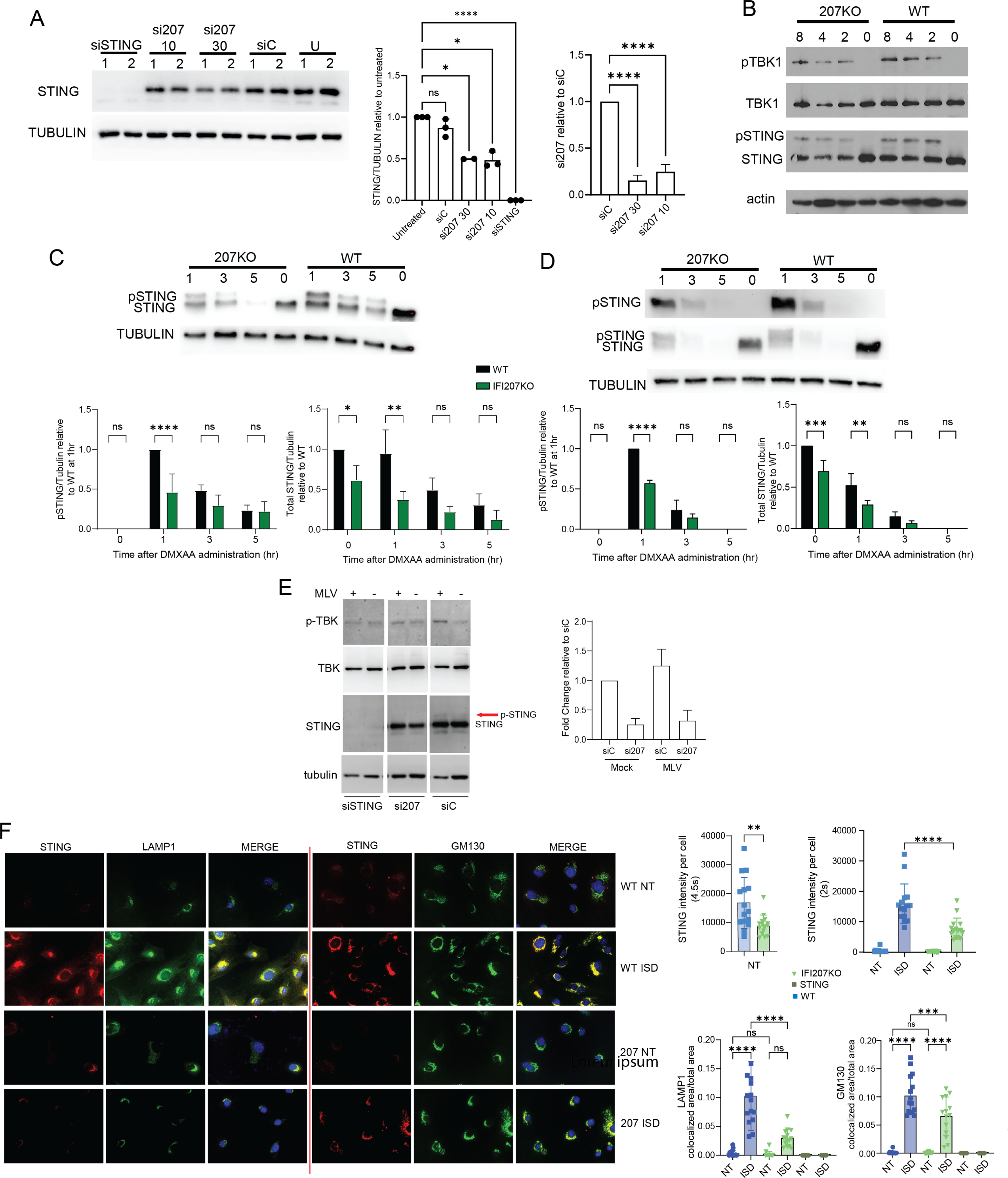
IFI207-deficient cells show decreased STING expression and diminished activation of the STING pathway. (A) NIH3T3 cells were transfected with the indicated siRNAs and western blots for endogenous STING were performed. Two concentrations of the 207 siRNA were used (10 and 30 pmole). Duplicate experimental replicates are shown. Quantification of western blots from 3 independent experiments and Ifi207 knockdown verification by RT-qPCR are shown to the right (average + SD). One-way ANOVA was used (*, *P*≤0.05; ****, *P*<0.0001; ns, not significant). Abbreviations: siC, control siRNA; U, untransfected. (B) BMDMs were transfected with ISD and at 2, 4 and 8 hr post-transfection, protein expression was analyzed by western blot with the indicated antibodies. Blots were stripped and probed with anti-actin to control loading. (C) and (D) Fibroblasts (C) and macrophages (D) from IFI207 KO and wild type mice were treated with 100μg/ml DMXAA and western blots examining STING and phospho-STING levels were measured at the indicated times. Shown below the graph is quantification of 3 independent experiments. Two-way ANOVA was used (*, *P*≤0.03; **, *P*≤0.001; ****, *P*<0.0001; ns, not significant). (E) NIH3T3 cells were transfected with siRNAs targeting IFI207 or STING. Non-targeting (siC) siRNA was also used. At 48 hr post transfection, cells were infected with MLV^gGag^ for 2 hr (MOI=5). Protein expression was examined by western blot with the indicated antibodies. Right panel: Ifi207 KD was verified by RT-qPCR. Average + SD from 2 independent experiments is shown. (F) Primary fibroblasts were transfected with 8μg /ml ISD and 2 hr post-transfection, the cells were fixed and stained with anti-STING and anti-LAMP1 (left) or anti-GM130 (right)). Shown to the right of the graphs is the quantification of STING staining from untreated cells (NT) and treated cells. Different exposure times were used to quantify STING levels in the absence (4.5 sec) and presence (2 sec) of ISD. Shown at the bottom are the fraction of cells with colocalized STING and LAMP1 or STING and GM130. G. 15 fields for each cell type/condition were analyzed. BMDCs from STING^mut^ mice are presented in Fig. S9B.

We then examined the levels of phospho-STING and TBK, the immediate downstream steps in the STING activation pathway. BMDMs from IFI207 KO and WT mice were transfected with ISD, and the presence of STING, phospho-STING, phospho-TBK1, and TBK1 were examined. The levels of total STING, phospho-STING and phospho-TBK1 were reduced in the IFI207 KO BMDMs compared to WT cells (Fig. 3B). Similar results were seen in ISD-transfected BMDCs (Fig. S8A). Finally, IFI207 KO fibroblasts and macrophages were treated with DMXAA, and STING and phospho-STING levels were analyzed by western blot. The KO cells had lower levels of both total and phospho-STING (Fig. 3C and 3D).

We also used siRNA knockdown to deplete *Ifi207* in NIH3T3 cells and infected them with MLV^gGag^, which because of its unstable capsid, falls apart in the cytoplasm more rapidly than wild type virus, and thus induces higher levels of IFNβ (18, 33). Knockdown of STING resulted in loss of phospho-TBK and -STING in response to infection (Fig. 3E).

Lower levels of endogenous STING protein in IFI207 KO fibroblasts were also seen by immunofluorescence microscopy (Fig. 3F); fibroblasts were used because of better visualization of the cytoplasm than other cell types. When STING is activated, it first moves to the Golgi and then to an acidic compartment, where it is degraded (36–38). To determine if this trafficking occurs in the absence of IFI207, we treated WT, IFI207 and STING^mut^ fibroblasts with ISD and stained them with the Golgi marker GM130 or LAMP1 (lysosomal compartment). STING staining in the absence of ISD was diffuse in both cell types in the absence of ISD (Fig. 3F). In contrast, ISD treatment caused STING to be localized in a perinuclear compartment in both cell types. No STING was detected in STING^mut^ cells (Fig. S9B). We also found that while there was less co-localization of STING in both compartments of IFI207 KO cells, the levels were particularly low in the LAMP1 positive compartment (Fig. 3F). Taken together, these data suggest in the absence of IFI207, lower levels of STING are present and may be the cause of the diminished anti-viral response.

### IFI207 binds STING

To better understand IFI207’s function and the possible reason for the S/T repeats expansion, we cloned the *Ifi207* coding regions from RNA isolated from the spleens of C57BL/6N, DBA/2J and a mouse from the 129P2/OlaHsD background (see Materials and methods), as representatives of the variants with different numbers of S/T repeats (Ifi207_BL/6_ = 12 repeats; Ifi207_DBA_ = 25 repeats; Ifi207_129_ = 24 repeats) (Fig. 1C and see below). For some experiments we also used molecular clones of *Ifi204* and I*fi203* derived from C57BL/6J mice for comparison (11).

To determine if IFI207 bound to STING, we co-transfected FLAG-tagged STING and all 3 full-length HA-tagged expression constructs (IFI207_BL/6_, IFI207_DBA_ and IFI207_129_) as well as an HA-tagged IFI204 vector and used anti-HA antibodies for immunoprecipitation and western blots. All 3 IFI207 constructs co-immunoprecipitated with STING (Fig. 4A). In contrast, although it was expressed at much higher levels than any of the IFI207 constructs, IFI204 only weakly co-immunoprecipitated with STING, (Fig. 4A).

**FIG 4.**
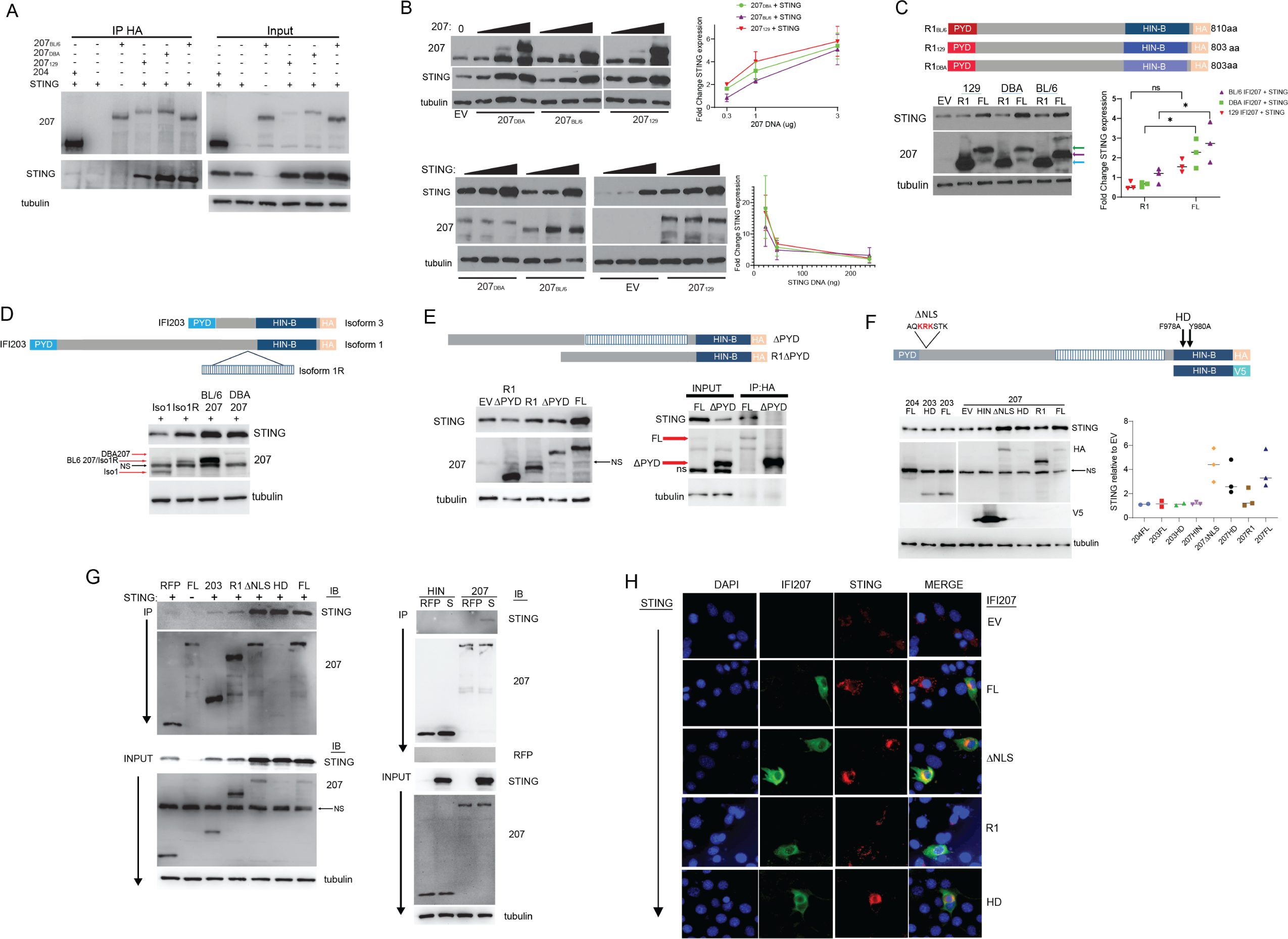
IFI207 stabilizes STING. (A) HEK293T cells were transfected with full length HA-tagged IFI207_129_ IFI207_BL/6_, IFI207_DBA_, IFI204 and FLAG-tagged STING expression constructs. Anti-HA was used to immunoprecipitate IFI207 in the transfected cell extracts, and western blots with the indicated anti-HA and -FLAG (STING) as well as anti-tubulin antibodies were performed. (B) HEK293T cells were transfected with increasing amounts of the indicated full length IFI207 (HA) (top) or STING (FLAG) (bottom) expression constructs. Protein expression was examined by western blot with antibodies to the tags and to STING. Experiments were done in triplicate and quantified by densitometry analysis in ImageJ (right panel). (C) HEK293T cells were co-transfected with HA-tagged full-length (FL) or repeat deletion (R1) IFI207_129_, IFI207_B6_, IFI207_DBA_ (diagram) and FLAG-tagged STING expression plasmids. Protein expression was examined by western blot with the antibodies to the tags and to STING. Experiments were done in triplicate and quantified by densitometry analysis in ImageJ (right panel). Green arrow, IFI207_129_, or IFI207_DBA_; purple arrow, IFI207_B6_; cyan arrow, R1 deletion constructs. (D) The same experiment was carried out with HA-tagged IFI203Iso1, IFI203Iso1R, IFI207_B6_ and IFI207_DBA_. (E) IFI207_DBA_ FL or R1 lacking the PYD domain were examined for their ability to increase STING levels (left panel) and coimmunoprecipitate with STING (right panel). (F) IFI207, IFI203 HD and IFI207 ΔNLS mutants and HIN constructs were tested for their ability to increase STING levels. All IFI207 plasmids are HA-tagged, except the HIN construct which contains a V5 tag. Protein expression was examined by western blot with the indicated antibodies; STING was detected with anti-STING antibody. Right panel: triplicate experiments were quantified by densitometry analysis in ImageLab (BioRad). Abbreviations: NS, non-specific; EV, empty vector. (G) HEK293T cells were co-transfected with the indicated IFI207, IFI203, IFI204 and FLAG-tagged STING expression plasmids. Immunoprecipitations were carried out with anti-HA (left panel) or V5 (right panel) and western blots were examined for co-immunoprecipitation with STING. (H) NIH3T3 cells were transfected with the full-length IFI207_BL/6_ (HA) with or without STING (FLAG) expression constructs. Cells were fixed and stained with antibodies directed against expression construct tags. Images were acquired by fluorescence microscopy. Image brightness was manually adjusted to better visualize STING in the STING-alone panels because of the lower levels of STING. Images of cells transfected with IFI207 constructs without STING are in Fig. S11B and quantification of subcellular localization in Fig. S11C.

Interestingly, we observed that even though equal amounts of STING plasmid were transfected in the presence or absence of IFI207, there was more STING protein in extracts when IFI207 was present (input; Fig. 4A). This was similar to our observation that endogenous STING protein levels were higher in IFI207-containing cells (Fig. 3). This suggested that STING stabilization might be dependent on IFI207/STING interaction. To determine if this was the case, increasing amounts of the IFI207 expression plasmids were co-transfected with an expression vector containing FLAG-tagged STING, and the levels of STING were examined by western blot. Co-expression of each of the IFI207 plasmids stabilized STING protein expression compared to empty vector (Fig. 4B). Moreover, both the IFI207_DBA_ and IFI207_129_ constructs stabilized STING more effectively than the IFI207_BL/6_ construct (Fig. 4B, top). Increasing amounts of STING plasmid were also co-transfected with constant amounts of the IFI207 plasmids, and again, all 3 IFI207 constructs increased STING protein expression. While IFI207 stabilized STING, increasing STING had no effect on IFI207 levels (Fig. 4B, bottom). This effect of IF207 was specific to STING; endogenous tubulin and actin expression was not affected by over-expression or depletion of IFI207 (Figs. 3 and 4) and expression of co-transfected FLAG-tagged RFP was not altered by co-transfection of IFI207 (Fig. S10A).

### The S/T-rich repeat region is important for STING stabilization

Although there are minor sequence differences in the PYD and HIN domains of the *Ifi207* genes found in BL/6, 129 and DBA mice, the major difference we found was in the sequence and number of the HIN-B-proximal S/T-rich repeat region; BL/6 IFI207 had only 12 repeats, while IFI207 in 129 mice lacked a single S/T repeat compared to DBA mice (24 vs. 25 repeats, respectively) but was otherwise identical (Fig. 1C; Tables S2 and S3). To assess whether the repeat region played a role in STING stabilization, we made constructs in the backbone of each allele that lacked all the S/T repeats (R1_BL/6_; R1_DBA_; R1_129_; Fig. 4C). These 3 constructs, as well as the full-length constructs from which they were derived (FL_BL/6_; FL_DBA_; FL_129_), were co-transfected into 293T cells with STING. Expression of all 3 R1 deletion constructs was much higher than the full length IFI207 constructs. Nonetheless, deletion of the repeat region abrogated the ability of IFI207 to stabilize STING, since the level of STING after co-transfection with R1_BL/6_, R1_DBA_ or R1_129_ was like that seen with empty vector (EV in Fig. 4C). Moreover, STING dimer formation increased with full-length but not the repeat deletion constructs (Fig. S10B).

To further confirm that the repeat region was critical for STING stabilization, we introduced the IFI207_BL/6_ HIN-B-proximal repeat region into the equivalent site in isoform 1 of IFI203 (Iso1) (RefSeq ID NP_001289578; see Fig. 5B). This isoform shares the same structure as the carboxy terminal half of IFI207 and is expressed more abundantly in primary macrophages than the other isoforms (12). The previously reported IFI203 expression vector contained the shortest isoform 3 (IFI203Iso3) (11). We introduced the IFI207_BL/6_ repeat region into IFI203Iso1 to make a chimeric IFI203 with 12 HIN-side S/T rich repeats (IFI203Iso1R; Fig. 1B). We co-transfected IFI203Iso1, IFI203Iso1R, FL-IFI207_BL/6_ and FL-IFI207_DBA_ with STING. IFI203Iso1R containing the BL/6 repeat region, but not IFI203Iso1 without the repeats, stabilized STING (Fig. 4D).

### STING stabilization does not rely on DNA binding or nuclear localization but does require the PYD domain

The PYD domain of IFI16 is required for its ability to interact with ASC, BRCA1 and STING (39, 40). We thus tested whether IFI207’s PYD domain was also required for STING stabilization. We deleted the PYD domain from both the FL-IFI207_DBA_ and R1-IFI207_DBA_ constructs and co-transfected them with STING. Only the full-length construct, but not the ΔPYD constructs stabilized STING (Fig. 4E). This suggests that although the repeat region was important for STING stabilization, the PYD domain was needed for STING binding. Indeed, the ΔPYD construct did not co-immunoprecipitate with STING (Fig. 4E).

In addition to the recent study suggesting that IFI207 enhances RNApolII binding at the transcription start sites of cytokine genes, it has been suggested that IFI16 restricts HIV-1 infection by interfering with the transcription factor Sp1, thereby suppressing transcription of the integrated provirus (17). To determine where IFI207 was located within the cell, we first performed fractionations of cells transfected with IFI207_BL/6_, IFI207_DBA_, IFI207_129_ expression vectors in the presence and absence of STING expression plasmids. While a large fraction of IFI207 was in the nucleus, it was also found in the cytoplasm, independent of the number of repeats; the BL/6, 129 and DBA constructs all showed both nuclear and cytoplasmic localization (Fig. S11A). STING also appeared in the nuclear fraction, likely due to its association with perinuclear membrane fractions (41, 42). Some of the IFI207 found in the nucleus may be due to a similar association.

We next examined three additional mutants for their ability to stabilize STING: ΔNLS, which deletes a putative nuclear localization signal downstream of the PYD; HD, with alanine substitutions at amino acids 978 and 980 in FL-IFI207_DBA_, which we showed previously abrogated nucleic acid binding when introduced into IFI203 (18); and HIN, which encodes only the HIN domain and was tagged with V5 (Fig. 4F). The ΔNLS and HD constructs showed decreased nuclear localization (Fig. S11B and C). We co-transfected expression constructs encoding these mutants as well as FL-IFI207_DBA_ and the R1 mutant with STING and examined STING stability by western blots. Constructs bearing IFI204, IFI203Iso3 and the DNA binding mutant IFI203 HD were also co-transfected with STING. We found that both IFI207 ΔNLS and HD mutants but not the HIN construct stabilized STING (Fig. 4F). Neither the IFI204 nor the IFI203 constructs stabilized STING.

To determine whether any of these regions also contributed to STING binding, we also carried out co-immunoprecipitations on extracts from cells co-transfected with STING and IFI207_DBA_ or the mutants on the DBA backbone. Although they were expressed at much lower levels than the R1 or HIN proteins, the FL, ΔNLS and HD mutants but not the R1 or HIN domain constructs efficiently bound and stabilized STING (Fig. 4G). As we showed previously, IFI203Iso3 also bound to STING, albeit less effectively than repeat-containing IFI207 and at levels similar to the R1 mutant (Fig. 4G) (18).

We also tested whether binding of reverse transcripts correlated with IFI207’s ability to stabilize STING. We showed previously that DDX41 and IFI203 but not the DNA binding mutant, IFI203 HD, precipitated MLV reverse transcription products from infected cells (18). MCAT cells (293T cells that stably express the MLV entry receptor ATCR1 (43)) were transfected with IFI207_DBA_-FL, -R1 and -HD, as well as IFI204, IFI203Iso3, and DDX41 and then infected with MLV^gGag^, which, because the capsid dissociates more rapidly than wild type virus, produces high levels of early reverse transcripts in the cytoplasm (18, 33). At 4 hr post-infection, the ability of the different constructs to precipitate MLV reverse transcripts was determined. IFI207_DBA_-FL and -R1 strongly bound MLV reverse transcripts, while IFI207 HD weakly bound viral DNA (Fig. S12). IFI204 and IFI203Iso3 also weakly bound viral DNA, as we previously showed (18). These data show that nucleic acid binding/activation is not required for IFI207 stabilization of STING, at least under conditions of over-expression.

Finally, we studied the co-localization of the different mutants with STING by immunofluorescence. Both the ΔNLS and HD mutants were more frequently localized in the cytoplasm, while like the WT protein, the R1 mutant was seen in both the nucleus and cytoplasm, both in the presence (Fig. 4H) and absence (Fig. S11B and C) of STING. STING immunostaining was brighter in the presence of IFI207, similar to what was seen in WT cells compared to IFI207 KO cells (Fig. 2). STING was also more often found in concentrated perinuclear foci in cells expressing the WT than the R1 constructs (Fig. 4H).

## DISCUSSION

Pathogens infect different cell types and tissues and many replicate in distinct cellular compartments. TLRs, ALRs, cGAS, and other sensors mediate innate immune responses to pathogens. The need for targeted responses to different pathogens has likely led to the expansion of PRRs with different signaling pathways. These specific sensing and signaling pathways are also important targets for pathogens, and as a result, greatly accelerate host-pathogen co-evolution. Indeed, in house mouse strains, *Alrs*, *Ifns* and even *Ifn* receptor genes are part of “strain-specific diverse regions” (SSDRs) with high sequence diversity (44). Some of the SSDRs are among the most dynamic regions in the genome. Both genome recombination (45), and non-crossover gene conversions (46) between large gene families with two divergent haplotype genomes will create novel gene structures. Unlike housekeeping genes, newly created immune-related genes have a higher chance of being beneficial and thus showing genetic drift in the population because of the “red queen hypothesis” (46). On the other hand, the destruction of an existing gene by recombination or gene conversion is more likely to be tolerated when it is part of a gene family because other family members may have (semi-) redundant functions. As a result, SSDRs are possibly also hot spots of new gene generation.

The *Alr* locus is a typical SSDR. With the exception of some bat species, it is found in all mammalian genomes (47). The ALRs belong to one family of innate immune sensors that may have diversified in response to the selective pressures of pathogens (10–13, 48). We show here that several new *Alr* family members have recently been introduced into the genomes of the *Mus* genus, probably by one or a series of unequal genome recombination events. Novel ALR members like MNDAL and IFI207 have acquired new roles in variable biology processes: MNDAL has become a cell growth regulator (49), and as we show here, IFI207 has acquired a key role in the STING mediated antiviral signaling pathway, using non-redundant and different mechanisms than the other ALRs. These new members may have modified the functional capacity of mouse ALRs, because in rats and humans, fewer ALR members are sufficient for normal immune function. For example, humans have *AIM2*, *IFI16*, *PYHIN1*, and *MNDA*, while rats have *Rn7* (*AIM2*), *Rhin2* (*Ifi206*), *Rhin3* (*Ifi214*), *Rhin4* (*Ifi203*) and *Rhin5* (*Mnda*) (Fig. 1B). Thus, no *Ifi207* orthologues are required in these species.

This kind of functional innovation by gene conversion or recombination may happen independently and repeatedly. In support of this is the structure of the *Ifi207*-like genes in *Apodemus sylvaticus*. During our analysis, we found more than 5 copies of novel genes with a structure similar to *Mus Ifi207* in this species. These genes are *A. sylvaticus* species-specific and lack the characteristic S/T rich repeats found in *Mus Ifi207*. However, they have undergone a recent gene duplication and seem to be under selection in other regions of the genes, resulting in highly polymorphic PYD-side repeats (Figs. S1 and S2). In addition, although these genes are possibly formed by recombination between *Ifi203* and *Mnda* as well, they seem to have different recombination sites from the *Mus* version. These results suggest that *Ifi207* in *Mus* and *A. sylvaticus* were introduced into genome independently and evolved to have different functions.

Although the entire *Alr* locus shows great diversity among different house mouse strains, *Ifi207* was strikingly diverse. We found that this gene contains a large HIN-B-proximal S/T-rich repeat region that varies extensively among inbred and wild mice strains. We analyzed the repeat region sequence in multiple mouse strains and found a large collection of different sequences of repeating units in the IFI207 repeat region. In addition to sequence variation of the repeat region, these mice also had different numbers of repeating units. Apart from this repeat region, the rest of the protein sequence was almost identical in different inbred strains, suggesting that the repeat region specifically is under strong selective pressure. There is a paradox between the young age (a few million years for the *Mus* genus, which may accumulate <2% mutation under neutral selection), and the extreme high variation (up to 36% at the nucleotide level) of the repeat units. The mechanism by which the sequence evolution occurs is yet to be resolved, but a better understanding of how the *Ifi*207 repeat region has evolved might offer clues about its role as an innate immune effector as wild mice encounter a wide array of pathogens that may have shaped its evolution.

The data presented here, using knockdown, knockout and over-expression approaches, show that IFI207 stabilizes STING protein and alters downstream signaling. The repeat region, which is predicted to form a coiled structure similar to a leucine zipper, makes IFI207 unique among the ALRs. The length of the repeat region also affected binding to STING and its stabilization. The IFI207 proteins encoded in DBA/2 and 129 mice, which have 25 and 24 repeats, respectively, stabilized STING more effectively than did the BL/6 protein, which has only 12 repeats. This suggests that these repeats expanded and were subsequently retained in the encoded protein, likely because of exposure to pathogens by the progenitor mice from which different inbred strains were derived. Several ALRs, including IFI16, IFI203, IFI204 and IFI205, bind STING, but these proteins lack the extended repeat region and as we show here, bind STING to the same extent as IFI207 with no repeats. In contrast, IFI203 engineered to contain the repeats increased STING protein levels. Although the PYD domain of IFI207, like other ALRs, confers binding to STING, it is the IFI207 repeat region that confers its ability to stabilize STING and likely enhances the response to different ligands and viruses that activate the STING/IFN response pathway.

A previous study suggested that knockout of the entire *Alr* locus in mouse macrophages had no effect on the response to ISD (50). However, a more recent study showed that these same ALR KO mice were more highly infected with MLV (17). A number of studies have implicated IFI204 in the response to herpesviruses, polyomavirus and several bacteria such as *Francisella* and *Staphyloccocus*, and IFI16 is important in human cells for the innate immune response to *Listeria*, herpesviruses and HIV (14, 17, 20, 51–53). We showed previously that IFI203, in conjunction with DDX41, recognized MLV and that DDX41 KO mice were as susceptible to MLV infection as cGAS KO mice (18, 21). We show here that IFI207 alters the response to DNA, cGAMP, DMXAA and MLV. Since MLV has been in *Mus* species for at least 1 million years, it is not surprising that there is an arsenal of restriction factors in mice that control MLV infection, including multiple ALRS (54). Whether IFI207 KO mice are more susceptible to infection with other pathogens that activate the STING pathway is currently under investigation.

Our results are somewhat at odds with a recent report by Baran and colleagues showing that IFI207 acts to increase RNApol II-mediated transcription of both NFκB and IRF3-stimulated genes (34). Both groups found that IFI207-depleted cells had impaired responses to DNA stimulation. However, unlike Baran et al., we found no significant effect of either LPS (TLR4 ligand) or bacteria treatment in a macrophage cell line depleted for IFI207 or primary BMDMs from IFI207 knockout mice. While the Baran paper demonstrated that IFI207-depleted cells had lower levels of TNFα and IFNβ1 in response to LPS or *K. pneumoniae*, IFI207 knockout mice in their study sustained lower levels of Klebsiella infection than WT mice. This is in contrast to infection of other mice with defects in the TLR or STING pathways, which are typically more highly infected by bacteria, and by our finding that IFI207 knockout mice are more susceptible to infection with MLV, consistent with the reduction in STING activation in the absence of IFI207.

IFI205 and IFI16 also alter gene expression, by acting as transcriptional repressors of ASC and integrated HIV proviruses, respectively, and these activities require nuclear localization (15, 17). Although IFI204 was shown to act as an anti-MLV and -HIV restriction factor by suppressing proviral transcription, we did not see any effect in IFI204 KO mice on infection levels, suggesting that it is not anti-viral *in vivo*. IFI207 also has an NLS, but we show that its ability to stabilize STING was independent of this sequence. Interestingly, at least in over-expression studies, the mutant lacking repeats still bound bind MLV reverse transcripts but failed to stabilize STING, while the HD mutant that has minimal nucleic acid binding still showed STING-stabilizing activity. The role of nuclear localization and the nucleic acid binding domain in IFI207 function thus remains to be determined.

There is incredible diversity in the *ALR* locus among mammalian species. For example, the horse locus has 6 *ALR* genes, dogs and pigs have 2, cows have 1, rats have 5, humans have 4 genes, and bats have only a truncated *AIM2* gene, all of which are encoded in a single locus which sits between the cell adhesion molecule 3 (*CADM3*) and spectrin alpha, erythrocytic 1 (*SPTA1*) genes (10–13, 47). While we show here that IFI207 enhances the anti-MLV and other STING-dependent innate immune responses, we do not know what pathogens may have shaped the diversity of this locus in different species. In addition to their roles in innate immunity, several ALRs have been implicated in adipogenesis (55, 56), autoimmune diseases like lupus erythematosus (57, 58) and inflammatory responses (15, 59). Identification of the pressures that shaped the locus in different species is thus important for understanding the many biological processes in which these genes play a role.

## MATERIALS AND METHODS

### Mice

C57BL/6N, DBA2/J, BALB/cJ, STING^mut^ (C57BL/6J-Sting1gt/J), AIM2 KO (B6.129P2-Aim2Gt(CSG445)Byg/J) and CMV-Cre (B6.C-Tg(CMV-cre)1Cgn/J) mice were originally purchased from The Jackson Laboratory and bred at UIC. IFI207 KO mice (C57BL/6N-IFI207em1(IMPC)Wtsi) were generated by targeted deletion by zygote injection of 4 gRNAs, two flanking each side of exon 3, to induce a frameshift null allele of *Ifi20*7. The embryos used for these experiments were C56BL/6NJ, and after germline transmission, the colony was maintained on this background. Full details of the gRNAs used for gene disruption are provided in Table S3A. IFI202 (Ifi202btm1(KOMP)Mbp/MbpMmucd) and IFI204 KO mice (Ifi204tm1a(KOMP)Wtsi/MbpMmucd) were derived from sperm purchased from the Mouse Biology Program and UC Davis. The IFI204 mice were initially crossed with CMV Cre mice to generate knockout strains and then backcrossed to remove the CMV Cre transgene (Fig. S4). Genotyping primers for the Ifi207, Ifi202 and Ifi204 knockout alleles are in Table S3B. cGAS KO mice (Mb21d1tm1d(EUCOMM)Hmgu) were originally provided by Michael Diamond and Skip Virgin (60). 129P2/OlaHsd DNA was isolated from mice originally purchased from Harlan (12).

*M. m. castaneus* mice were purchased from the Jackson Laboratory. All mice were housed according to the policies of the Animal Care Committee (ACC) of the University of Illinois at Chicago (UIC); all studies were performed in accordance with the recommendations in the Guide for the Care and Use of Laboratory Animals of the National Institutes of Health. The experiments performed with mice in this study were approved by the UIC ACC (protocol no. 21-125).

### Sequence analysis

The assemblies of non-mouse species are downloaded from NCBI (https://www.ncbi.nlm.nih.gov), with access names listed in Table S1 and S2. Whole genome de novo assembly, as well as the raw PacBio / PacBio Hifi reads of 13 mouse strains, *Mus spretus* and *Mus caroli* were provided by the Phase 3 Mouse Genomes Project (MGP) from EMBL-EBI (unpublished data; available from NCBI/EBI under project accession: PRJEB47108). The assemblies were produced using third generation long reads (PacBio Sequel / PacBio Hifi, BioNano®) and Hi-C. For the additional mouse strains, the alleles of Ifi207 were assembled from whole genome Illumina data or the transcriptomes of IFNα/γ-induced diaphragm fibroblast cells. Samples were provided by Institute of Vertebrate Biology CAS. The sequences of HIN-B-proximal S/T repeat regions were double-confirmed by Sanger sequencing of genome amplicons, with the primers described in Table S7. Genomic DNA samples from mouse strain SIN, SIT, BULS, PERC, BULS, STUF, DJO, MOLF and CIM were acquired from a variety of sources, and stored in Gulbenkian Institute of Science, Portugal. The details of mouse strains can be found in website https://housemice.cz/en/strains/. Full-length sequences of DBA and 129 IFI207 were obtained from the cloned cDNAs (see Plasmids and Transfection). Genbank acquisition numbers for the re-sequenced repeat regions and full-length clones are listed in Supplementary Table S5. For more details, see Plasmids and Transfection.

### Cells

NIH3T3 and HEK293T cells were cultured in Dulbecco’s Modified Eagle Medium (DMEM) supplemented with 10% FBS, 2mM L-glutamine (Gibco), 100 U/ml penicillin (Gibco), and 100 mg/ml streptomycin (Gibco). Cell cultures were maintained at 37°C with 5% CO2. MCAT cells were cultured in DMEM supplemented with 8% donor bovine serum, L-glutamine, penicillin/ streptomycin and G418 (10 mg/ml) (Goldbio). Mouse macrophage cell line NR9456 was cultivated in Dulbecco’s modified Eagle medium (DMEM; Invitrogen) supplemented with 5% fetal bovine serum (FBS; Invitrogen), glutamine (2 mM), sodium pyruvate (1 mM; Mediatech) and penicillin (100 U/ml)-streptomycin (100 µg/ml) (Invitrogen). Fibroblasts were harvested from tails or ears. The tissues were treated with collagenase (1000 unit/ml in Hepes Buffered Saline) and fibroblasts were cultured in DMEM containing 10% FBS, 2 mM L-glutamine, 100 U/ml penicillin, 100 ug/ml streptomycin at 37°C with 5% CO2.

### BMDM and BMDC cultures

Bone marrow was harvested from mouse limbs. To differentiate BMDMs, cells were cultured for 6-7 days on Petri dishes in DMEM containing 10% FBS, 2 mM L-glutamine, 100 U/ml penicillin, 100 μg/ml streptomycin, 0.1 mM sodium pyruvate, 25mM HEPES (Gibco), and 10 ng/ml murine M-CSF (Gibco, PMC2044). Fresh media containing 10 ng/ml M-CSF was added on day 3. On day 6 or 7, BMDMs were collected for experiments. To differentiate BMDCs, bone marrow cells were treated with ACK, and 2×10^6^ cells were seeded in the center of Petri dishes in 10ml RPMI containing 10% FBS, 2 mM L-glutamine, 100 U/ml penicillin, 100 μg/ml streptomycin and 50 μM β-mercaptoethanol plus 20 ng/ml murine granulocyte-macrophage colony-stimulating factor (GM-CSF) (Peprotech, 315-03). On day 4, 10 ml fresh media containing 20 ng/ml GM-CSF was added to plates. On day 8, half of the culture volume was carefully removed to avoid disrupting the cells clustered in the centers of dishes, and 10 ml fresh media with 20 ng/ml GM-CSF was added to plates. Non-adherent cells clustered in centers of wells were collected by gentle pipetting for experiments on day 10. At time of collection, BMDMs and BMDCs were stained for F4/80, CD11b, or CD11c and analyzed by flow cytometry (LSR-Fortessa flow cytometer; BD Biosciences). Macrophage and DC cell populations were >80% pure.

### Expression Analysis

BMDMs and BMDCs from BL/6, STING^mut^, IFI202, IFI204 and IFI207 KO mice were treated with or without 100 U mouse IFNβ (PBL Assay Science, 12405-1). Four hr after treatment, RNA was harvested, and Alr gene expression was analyzed by RT-qPCR. RNA was isolated by RNeasy Mini Kit (QIAGEN 74106) with on-column DNase digestion using RNase-Free DNase Set (QIAGEN 79254). cDNA was synthesized with random hexamers from the Superscript III First-Strand Synthesis System (Invitrogen 18080051). RT-qPCR was performed using Power SYBR Green PCR Master Mix (Applied Biosystems 4367659) with appropriate primer sets on a QuantStudio 5 Real-Time PCR System (Applied Biosystems). Primers for RT-qPCR analysis of *Alrs*, *Ifnb*, *Actin*, and *Gapdh* have been previously described (Table S5) (12, 18). MLV strong-stop primers have been previously described (61).

### ISD, cGAMP and DMXAA stimulation of BMDMs, BMDCs and fibroblasts

BMDMs were transfected with 8, 4, 2, 1, or 0 µg/ml interferon stimulatory DNA (ISD) (InvivoGen) or 2’3’-cGAMP (Sigma) for 3 hr. Fibroblasts were transfected with 8μg /ml of ISD or cGAMP and harvested after 3 hr. Transfections were done using Lipofectamine 3000 (Invitrogen. For western blot analysis, cells were transfected with ISD for 2 h. After 2 h, culture media was removed and replaced with fresh media, and samples were harvested at the time points indicated in the figure legends. For DMXAA (Sigma Aldrich, D5817) treatment, cells were treated without or with DMXAA at 100 µg/ml for 1, 3 and 5 hr. After stimulation, RNA was isolated and IFNβ mRNA expression was analyzed by RT-qPCR. For western blots, the cells were collected in PBS, pelleted, lysed in radioimmunoprecipitation (RIPA) buffer (50mM Tris pH 8, 150mM NaCl, 1mM EDTA, 1% Triton X-100, 1% sodium deoxycholate, and 0.1% SDS), and prepared as described in the western blot section below.

### siRNA knockdowns

NR9456 and NIH3T3 cells were reverse transfected with 30pmol of siRNAs (Silencer Select, Ambion by Life Technologies) targeting Ifi207 (s105479, s105481, s202438), using Lipofectamine RNAiMAX (Invitrogen). Non-targeting control siRNA was also included (AM4635). Knockdown efficiency was determined by RT-qPCR with primers specific for each target gene. The selectivity of the siRNAs for IFI207 was determined by RT-qPCR (Fig. S7A).

### LPS stimulation of BMDMs and macrophages

NR9456 were reverse transfected with 30pmol of IFI207 or control siRNAs. 48 hr post-transfection, the cells were treated with 10ng/ml LPS (Lipopolysaccharides from Escherichia coli O55:B5; Sigma). After 3 hr, RNA was collected, and *Tnf* and *Ifnβ1* RNA levels were analyzed by RT-qPCR. Primary BMDMs were treated with 10ng/ml LPS and RNA for RT-qPCR analysis was isolated at 3 and 6 hr post-treatment.

### Bacterial infection of BMDMs

For *S. aureus* (USA300 LAC), overnight bacterial cultures were added into a new tube containing fresh tryptic soy broth (TSB) and incubated for 3 hr at 37℃. Bacteria were pelleted, washed twice with PBS and adjusted to OD600 of 1.0 (1×10^9^ CFU/ml). BMDMs from WT, IFI207KO, STING^mut^ and IFI204 KO mice were infected by *S. aureus* at multiplicity of infection (MOI) of 5 and 100. Gentamicin was added 1 hr after infection, the media with bacteria was removed, the cells were washed twice with PBS and fresh media with 200 µg/ml of gentamicin was added. Four hr after infection, RNA was collected, and *Tnf*/*Ifnβ1* mRNA expression was analyzed by RT-qPCR. For *K. pneumoniae* (KPPR1), overnight bacterial cultures were grown in LB broth and a 1:10 dilution was inoculated into fresh LB and incubated for 2.5 hr at 37℃. Bacteria were pelleted and adjusted to an OD 600 of 1.0 (1×10^10^ CFU/ml). BMDMs and BMDCs from WT, IFI207KO, STING^mut^ and IFI204 KO mice were infected with *K. pneumoniae* at a MOI of 10 and 100. At 1 hr after infection, the media with bacteria was removed, the cells were washed twice with PBS, and fresh media with 100µg/ml of gentamicin was added. At 6 hr after infection, cells were harvested for RNA isolation.

### Virus

Wild-type Moloney MLV and MLV^gGag^ were harvested from supernatants of stably infected NIH3T3 cells. Supernatants were centrifuged for 10 mins (3000rpm) to remove cellular debris, and then filtered through a 0.45 µm filter (Millex Millipore) and treated for 30 min at 37°C with 20 U/ml DNase I (Roche). The virions were pelleted through a 25% sucrose cushion, resuspended in 2% FBS-supplemented DMEM, snap frozen and stored at -80°C. Virus titers were determined by an infectious center (IC) assay on NIH3T3s. One day before titering, 8×10^4^ cells were plated in 6-well tissue culture plates and incubated overnight at 37°C. Ten-fold serial dilutions of virus stocks were prepared with polybrene (8 µg/ml) in supplemented DMEM and added to cells. Infected cells were incubated at 37°C for 2 hr with rocking every 15 min. After 2 h, viral dilutions were aspirated, and 2.5 ml media was added to each well. Plates were incubated for 3 days at 37°C. After 3 days, plates were washed in PBS and stained with monoclonal 538 Env antibody (1:300) for 1 hr at 4°C with rocking. The plates were washed with 2% FBS in PBS and stained with Goat anti-mouse IgG/IgM Alexa Fluor 488 conjugated secondary antibody (Invitrogen A-10680) diluted in 2% FBS in PBS (1:300) for 45 mins at 4°C with rocking. Plates were washed and ICs were counted by microscopy (Keyence BZ-X710 and BZ-X analyzer) to determine viral titers. For virus stocks derived from cells transfected with molecular clones, viral mRNA was quantified by RT-qPCR with primers for env and viral proteins were confirmed by western blotting with MLV antisera, as previously described (31).

### *In vitro* MLV infection

Forty-eight hr post siRNA transfection, cells were infected with MLV^gGag^ and harvested at 2 hr post infection. Cells were collected and lysates were analyzed by western blot.

### *In vivo* MLV infections

IFI207 heterozygous mice were crossed, and their newborn litters consisting of knockouts and heterozygotes, as well as WT mice, were used for infection. Genotyping was done after virus loads were measured. Additional BL/6N pups were also infected and included in the WT group. STING^mut^ and IFI204 KO mice were generated by homozygote matings. Mice were intraperitoneally injected with 2×10^3^ PFU MLV at 0-48hrs-post birth. At 16 days post-infection, splenocytes were harvested and treated with ACK Buffer for RBC lysis. Tenfold dilutions of the cells were co-cultured with NIH3T3 cells for 3 days to assess viral titers by IC assays.

### *In vivo* LPS treatment

WT, IFI207KO, TLR4 KO and IFI204 KO mice were injected into the right hind footpad of with 1μg of LPS. At 2 hr after LPS injection, cells were isolated from the popliteal draining (right) and non-draining (left) lymph nodes. RNA was isolated, and *Tnf* and *Ifnβ1* RNA levels were analyzed by RT-qPCR.

### Plasmids

Expression plasmids for DBA and 129 IFI207 variants were generated by PCR amplification of cDNA synthesized from splenic RNA of DBA2/J and AIM2 KO mice. AIM2 KO were used because 129/P2 mice were no longer available for purchase, and the Alr locus, and *Ifi207* in AIM2 KO mice is derived from the parental 129P2/OlaHsd embryonic stem cells in which the knockouts were made (12). An HA tag was added to the C-termini immediately before the stop codon of both IFI207 variants. The PCR products were cloned into pcDNA3.1/Myc-His(+) (Invitrogen). Full-length sequences of DBA and 129 IFI207 were obtained from the cloned cDNAs. IFI207 R1 mutants, 𝛥NLS, ΔPYD and HD mutants were generated from full length IFI207-HA plasmids and the R1-ΔPYD mutant from R1-IFI207 using the Q5 Site-Directed Mutagenesis Kit (New England Biolabs). IFI207-HIN plasmids were generated by PCR amplification of the HIN domain from full length IFI207-V5 expression plasmids. BL/6-derived IFI203Iso2-HA, IFI207_BL/6_-HA, IFI204-HA and STING-FLAG plasmids were a gift from Dan Stetson (11). The IFI203Iso2 HD plasmid was previously described (18). V5 tags were introduced to the IFI207_BL/6_, IFI207_DBA_ and IFI207_129_ constructs by PCR amplification of the IFI207-HA expression constructs with primers designed to replace the HA tags with V5 tags. The V5-tagged PCR products were cloned into pcDNA 3.1/Myc-His(+).

The IFI203Iso1-HA plasmid was generated by in-fusion cloning. In brief, cDNA from C57BL/6N primary fibroblasts was used as a template to amplify full-length IFI203 isoform 1, using primers 203F and 203R. PCR was performed to linearize IFI203Iso2-HA plasmid using primers B6 203 1F and B6 203 2R. The insertion DNA fragment from the isoform 1 cDNA was PCR-amplified using primers B6 203 3F and B6 203 4R. In-fusion assembly with the 2 fragments was performed using the In-fusion snap assembly kit by Takara Bio.

To generate IFI203Iso1R containing the IFI207 repeat region, PCR was performed to linearize IFI203Iso1-HA plasmid using primers 203 VF and 203 VB. The repeat region of IFI207_BL/6_ was amplified by PCR using primers 207 21bpF and b6 21bpR. After purification, the PCR products were used for in-fusion assembly.

All cloning primers are described in Tables S6 and s7. The myc-tagged DDX41 construct was previously described (18). All plasmids were sequenced to verify the inserts.

### Transfections for Protein Expression Analysis

HEK293T cells were transiently transfected using Lipofectamine 3000 for 24 h. For fractionation assays, HEK293T cells were transfected with expression plasmids, collected in ice cold PBS, and fractionated into nuclear, cytoplasmic, and whole cell extracts by the Rapid, Efficient And Practical (REAP) method (62). The REAP method was modified to lyse nuclear extracts in RIPA buffer and to treat nuclear lysates with 3U Micrococcal Nuclease (Thermo Scientific) per µl lysate for 15 min at RT. For IFI207 and STING titration assays, HEK293T cells were transfected with either increasing amounts of IFI207-HA and a constant amount of STING-FLAG or with increasing amounts of STING-FLAG and a constant amount of IFI207-HA. When different amounts of plasmids were transfected within experiments, empty vector (EV) plasmid (pcDNA 3.1/Myc-His(+)) was added to keep the total amount of transfected DNA equal for all samples.

### Immunoprecipitations

For immunoprecipitation (IP) assays, HEK293T cells were transfected with expression plasmids for 24 hr using Lipofectamine 3000. Cells were harvested in PBS, pelleted, lysed in RIPA buffer, sonicated, and quantified by Bradford Assay (Bio-Rad). IP samples were prepared with Protein A/G PLUS-Agarose (Santa Cruz sc-2003) or HA-tag and V5-tag Sepharose bead conjugate (3956S and 67476S, respectively). Antibodies and Agarose or Sepharose beads were incubated with cell lysates at 4°C overnight on a rotating rack. Samples were washed 3-5 times in RIPA buffer and centrifuged 3 min (2500 rpm). Samples were eluted in 4X Laemmli buffer plus 5% β-mercaptoethanol.

### Western blots

For all protein expression analysis, lysates were sonicated on ice, quantified by Bradford Assay, prepared in 4X Laemmli buffer plus 10% β-mercaptoethanol, and boiled for 5 mins. Protease and Phosphatase inhibitor cocktail (Thermo Scientific 78443) was added to all lysis buffers immediately before use. Proteins from cell lysates were resolved by sodium dodecyl sulfate polyacrylamide gel electrophoresis (SDS-PAGE) and then transferred to polyvinylidene difluoride membranes (Millipore IPVH00010). For native blots, lysates were prepared in 4X loading buffer without SDS, and gels were run with Tris-glycine running buffer without SDS. Proteins were detected with primary antibodies against HA-tag (Cell Signaling Technology (CST) C29F4 or Abcam 9110), FLAG-tag (Sigma F1804 or CST 2368), V5-tag (Invitrogen 46-0705), STING (CST D2P2F or D1V5L), lamin B1 (CST D4Q4Z), ⍺-tubulin (Sigma T6199), TBK1 (CST 3013), and phospho-TBK1 (Ser172) (CST D52C2). HRP-conjugated antibodies (CST anti-rabbit or anti-mouse HRP IgG,7074 and 7076, respectively) were used for detection with Pierce ECL Western Blotting Substrate (Thermo Fisher Scientific, 322209) or ECL Prime Western Blotting Reagent (GE Healthcare Amersham, RPN2232).

### Indirect Fluorescent Antibody (IFA) Assay

NIH3T3 cells were seeded on glass coverslips in 12-well plates and incubated at 37°C overnight. Cells were transfected with IFI207-HA and STING-FLAG expression plasmids for 24 hr with Lipofectamine 3000. To stain, the culture media was aspirated, the cells were fixed in 4% paraformaldehyde (PFA) for 10 min, washed in PBS for 3 x 2 min, permeabilized in 0.25% Triton X-100-PBS for 5 min, washed, blocked for 30 min in 10% goat serum/PBS, incubated in primary antibody 2 h, washed, incubated in secondary antibody 2 h, and washed a final time. Coverslips were mounted on slides with VECTASHIELD Antifade Mounting Medium with DAPI (Vector Laboratories, H-1200). Primary and secondary antibodies were diluted in blocking buffer. Primary antibodies used were rabbit anti-HA (Abcam 9110) and mouse anti-FLAG M2 (Sigma, F1804). Goat anti-Rabbit IgG Alexa Flour 568-conjugated (Invitrogen, A-11011) and goat anti-mouse IgG Alexa Fluor 488-conjugated (Invitrogen, A-11001) secondary antibodies were used. Images were captured on a Keyence BZ-X710 at 60X magnification under oil immersion. Image brightness was manually adjusted to better visualize STING. Scoring of images was done in a blinded fashion. Images were scored for IFI207 and STING localization (cytoplasmic, nuclear, or both). To quantify localization, cells were counted that were IFI207-positive or STING-positive (single transfections) or IFI207- and STING-double positive (double transfections). Cells scored as cytoplasmic, nuclear, or both are represented as the fraction of the total cells counted within each transfection group (Fig. S11B).

Primary fibroblasts were seeded on glass coverslips in 24-well plates and incubated at 37°C overnight. ISD was transfected into fibroblasts for 2 hr with Lipofectamine 3000 reagent (Invitrogen). Fibroblasts were prepared for staining as described above. The fibroblasts were incubated with primary antibody overnight at 4°C and then stained with the secondary antibody (all 1:1000, Alexa Fluor; Invitrogen) for 1 h. Coverslips were mounted on slides with VECTASHIELD Antifade Mounting Medium with DAPI (Vector Laboratories, H-1200). The antibodies and concentrations used were as followed: sheep anti-STING (1:100, R and D Systems AF6516, RRID:AB_10718401), rabbit polyclonal anti-GM130 (2ug/ml, Novus NBP2-53420G, RRID:AB_2890912) and rabbit anti-LAMP1 (1:100, Sigma-Aldrich L1418, RRID:AB_477157). Images were acquired using a Keyence BZ-X710 at 40X and 60X magnification. Data analysis was performed with BZ-X Analyzer software. STING intensity analysis was calculated by averaging the STING fluorescence intensity per total number of cells.

### Nucleic acid pulldowns

Pulldowns were performed as previously described (20). Briefly, 293T MCAT cells transfected with pcDNA3.1 (empty vector) or the indicated expression constructs were infected with virus and at 4 hpi, the cells were cross-linked with 1% formaldehyde in media. Cross-linking was quenched with 2.5M glycine, and then extracts were incubated overnight with anti-myc- or anti-HA-agarose beads (Sigma) and G/A-agarose beads (SantaCruz). The beads were washed with high-salt buffer (25mM Tris-HCl, pH 7.8, 500mM NaCl, 1mM EDTA, 0.1% SDS, 1% TritonX-100, 10% glycerol) and with LiCl buffer (25mM Tris-HCl, pH 7.8, 250mM LiCl, 0.5% NP-40, 0.5% Na-deoxycholate, 1mM EDTA, 10% glycerol). The immunoprecipitated nucleic acid was eluted from the beads at 37°C in 100mM Tris-HCl, pH 7.8, 10mM EDTA, 1% SDS for 15 min, and the protein-nucleic acid cross-linking was reversed by overnight incubation at 65°C with 5M NaCl. The eluted nucleic acid was purified using the DNeasy Kit (Qiagen) and analyzed with RT-PCR strong stop primers (SSS) (Table S5) (61).

### Statistical Analysis

All gene expression RT-qPCR analysis experiments were done in duplicate or triplicate with 2-3 technical replicates for each experiment. Each technical replicate was additionally split into triplicates for RT-qPCR analysis. All other experiments were performed at least in triplicate with technical replicates or as indicated in the figure legends. Fluorescence quantifications were done in a blinded fashion. GraphPad Prism was used for statistical analysis. Tests used to determine significance are shown in the figure legends.

## ACKNOWLEDGEMENTS

We thank David Ryan for help with the mouse breeding and analyzing the immunofluorescence data, and the members of our lab for helpful discussions.

## FUNDING

Supported by National Institutes of Health Grants R01AI121275 and 1R01AI174538 (SRR).

## AUTHOR CONTRIBUTIONS

Conceptualization: JL, SRR

Methodology: JL

Investigation: EM, KSB, WZ, TE, ANA, TMK, JL, IA, CA, DK

Supervision: SRR, FA, JB

Writing—original draft: EM, KSB, WZ, TE, JL, SRR

Writing—review & editing: EM, KSB, WZ, TE, JB, DK, FA, IA, DJA, JL, SRR

## COMPETING INTERESTS

Authors declare that they have no competing interests.

## DATA AND MATERIALS AVAILABILITY

Raw data, including all cell images scored in Figs. 2, 3, S7 and S11, are available in the Mendeley file “IFI207”, Mendeley Data, V1, doi: 10.17632/d922×2wzf8.1. Whole genome sequences for the mouse strains are available from NCBI/EBI under project accession: PRJEB47108.

